# Effects of one-night partial sleep deprivation on perivascular space volume fraction: Findings from the Stockholm Sleepy Brain Study

**DOI:** 10.1101/2024.10.26.620382

**Authors:** Rachel M. Custer, Kirsten M. Lynch, Giuseppe Barisano, Megan M. Herting, Torbjörn Åkerstedt, Gustav Nilsonne, Hedyeh Ahmadi, Jeiran Choupan

**Author notes:** **Corresponding Author:** Jeiran Choupan, Laboratory of Neuro Imaging (LONI), Mark and Mary Stevens Neuroimaging and Informatics Institute, Keck School of Medicine, University of Southern California, Los Angeles, CA, USA.

## Abstract

Increased waste clearance in the brain is thought to occur most readily during late-stage sleep (stage N3). Sleep deprivation disrupts time spent in deeper sleep stages, fragmenting the clearance process. Here, we have utilized the publicly available Stockholm Sleepy Brain Study to investigate whether various sleep-related measures are associated with changes in perivascular space (PVS) volume fraction following a late-night short-sleep experiment. Our sample consisted of 60 participants divided into old (65-75 years) and young (20-30 years) age groups. We found that partial sleep deprivation was not significantly associated with major PVS changes. In the centrum semiovale, we observed an interaction between percentage of total sleep time spent in N3 and sleep deprivation status on PVS volume fraction. In the basal ganglia, we saw an interaction between N2 (both percentage of total sleep time and absolute time in minutes) and sleep deprivation status. However, the significance of these findings did not survive multiple comparisons corrections. This work highlights the need for future longitudinal studies of PVS and sleep, allowing for quantification of within-subject morphological changes occurring in PVS due to patterns of poor sleep. Our findings here provide insight on the impacts that a single night of late-night short-sleep has on the perivascular waste clearance system.

**Statement of Significance:** Sleep issues have been known to be associated with a variety of health risks. Recent research has shown the importance of sleep for brain clearance, with increases in brain clearance occurring during sleep. PVS seen on MRI are known to represent both influx and efflux pathways for the exchange of cerebrospinal fluid (CSF) and interstitial fluid (ISF) in the brain parenchyma, a key process underlying the waste clearance system.^1^ Numerous recent studies have found links between sleep disruption and PVS volume changes. However, no study to date has been able to examine within-subject changes in PVS resulting from experimental manipulation of sleep. Our study represents the first within-subject study looking at the effects of partial sleep deprivation on PVS volume fraction using an automated PVS segmentation technique.

## Introduction

Sleep deprivation is known to have a multitude of negative effects on both the human brain and body. Studies suggest that sleep deprivation causes decreases in pain tolerance, increases in circulatory pro-inflammatory biomarkers, and changes in social decision making.^2–4^ Partial sleep deprivation, defined as less than six hours of sleep per night,^5^ has been shown to affect cognition, impairing working memory and attention.^6^ Moreover, studies examining the effects of partial sleep deprivation on immune functioning have found that sleep deprivation can cause changes in T cell cytokine production and natural killer cell responses, potentially contributing to future pathologies and immune deficiencies.^7^

Sleep can be divided into several stages within two main categories - rapid eye movement (REM) sleep and non-rapid eye movement sleep (NREM).^8^ REM consists of a single stage alone, while NREM can be further subdivided into the stages of N1, N2, and N3. In a single night of sleep, one progresses from the lighter N1 stage through to the subsequently deeper stages of N2 and N3 sleep, before returning to N2 and then progressing to REM. This constitutes a single cycle of sleep and lasts approximately 90-110 minutes.^8^ Most sleep takes place in NREM, but proportion of time spent in REM sleep increases as time spent sleeping increases.

Waste clearance in the brain occurs most readily during sleep,^9^ and particularly during N3 (slow-wave sleep). This is due to the large brain wave oscillations present during N3 sleep, which cause increased CSF flow into the interstitial space.^10^ Such influx of CSF into the interstitial space facilitates waste clearance within the brain, which is carried through fluid filled spaces surrounding blood vessels called PVS.^11,12^ Research suggests that as part of the brain clearance system, PVS work to flush harmful waste products like beta-amyloid through the interstitial space.^13^ While much of the waste clearance is currently hypothesized to occur during N3 sleep, the presence of occasional delta waves during N2 sleep suggest that other stages may also have beneficial and contributory roles. However, current understanding suggests that N1 (superficial sleep) and rapid-eye-movement sleep (REM), though possibly involved in sleep-related PVS changes, are unlikely to be the primary drivers of these changes.^14,15^

High-resolution imaging performed at 17.6T on anesthetized rats has allowed for further description of the pathways involved in the brain’s PVS network, showing connections between ventricles and brain parenchyma and likely representing the overall path of solute clearance.^16^ PVS may become enlarged due to disruptions in this brain clearance system, whereby CSF may accumulate inside of the PVS and cause them to become visible on T2-weighted MRI as hyperintense tubular structures within the brain’s white matter.^17^ Enlarged PVS were initially thought to be a benign finding, as enlargement typically occurs with aging.^13^ Recent research however, has suggested that enlargement may signal underlying processes that accompany cerebral small vessel disease and other pathologies.^17^ Enlarged PVS have been associated with depression, diabetes, and Alzheimer’s disease in the elderly population.^18^ Increased prevalence of enlarged PVS in the centrum semiovale (CSO) of the brain’s white matter have also been noted in those with early cognitive dysfunction as well as those with hypertension.^19^ Studies examining brain changes involved in neurodegenerative disorders, such as Parkinson’s disease (PD), identify increases in both global and regional PVS volume fractions in those with PD when compared to controls.^20^ Thus, changes in PVS and the underlying deficits in waste clearance that likely precede such changes appear to be linked to a multitude of pathologies and risk factors for cognitive decline.

Links between the proportion of time spent in various sleep stages and enlarged PVS have been found in several recent studies. One study found that higher sleep efficiency measured via actigraphy was linked to higher PVS burdens in the CSO.^21^ Alternatively, another study found that the proportion of participants with poor sleep quality was higher in groups with large quantities of PVS in the basal ganglia (BG-PVS) compared to a control group.^22^ They found the same relationship in participants with large amounts of PVS in the CSO compared to controls.^22^ An additional study investigated the relationship between time spent in each sleep stage and enlarged PVS.^14^ Here, they found that more time spent in the N1 stage of sleep (measured in both absolute and proportional terms) and a lesser duration of absolute N3 sleep were associated with enlarged PVS.^14^ Recently, it was reported that there are differences in the relationship between sleep and PVS depending on age, with sleep efficiency being negatively associated with basal ganglia PVS (BG-PVS) volume fraction in older adults, and positively associated with BG-PVS in middle-aged adults.^23^ This suggests that the mechanisms underlying PVS enlargement may change with age, encouraging investigations into how these differences manifest in each group independently. Limitations of the previous studies involve lack of experimental manipulation of sleep to observe if acute disruptions in sleep also affect PVS burden. Here, we investigate whether one single night of partial sleep deprivation results in observable changes in PVS volume fraction within-subject measured via MRI. Using the Stockholm Sleepy Brain 1 dataset (which involves a late-night short-sleep design)^24,25,26^ we have investigated how sleep deprivation relates to changes in CSO-PVS and BG-PVS volume fraction in relation to changes in various polysomnography-derived measures, such as changes in proportion and duration of time spent in N1, N2, N3, REM, and the combined measure of N2+N3.

Since methods that rely upon automated quantification of PVS volumes have a greater ability to be sensitive to miniscule changes in global PVS measures, we have used an automated PVS segmentation technique developed by our team.^27,28^ Additionally, we explored how changes in PVS are associated with alterations in sleep stages, as assessed with polysomnography recordings. We expect such changes to be associated with decreases in total sleep time, time spent in deep sleep, and sleep efficiency. This study allows us to examine common sleep-related factors that may drive consistent MRI-visible PVS change both within and across subjects.

## Methods

### Sample

Data was obtained from the openly available Stockholm Sleepy Brain Study 1.^25^ Details about the study, including inclusion/exclusion criteria, participant recruitment, and detailed analysis and data collection methods can be found in the open-access data archive and the original study manuscripts.^24,25^ However, in brief, T1w images were acquired following a normal night of sleep and a night of sleep deprivation (where participants were instructed to sleep only 3 hours), which were counterbalanced so that approximately half of the participants received the sleep deprivation condition first and the other half received the night of normal sleep first. These scans happened at an interval of approximately one month apart (**Figure 1**). T2w FLAIR images were also acquired at the first scan session for all participants, regardless of whether they were in the sleep deprived or normal sleep condition at that time point. Polysomnography was acquired during the night preceding the scan on each of these separate occasions, and sleep recording took place in the subjects’ own homes.

**Figure 1:**
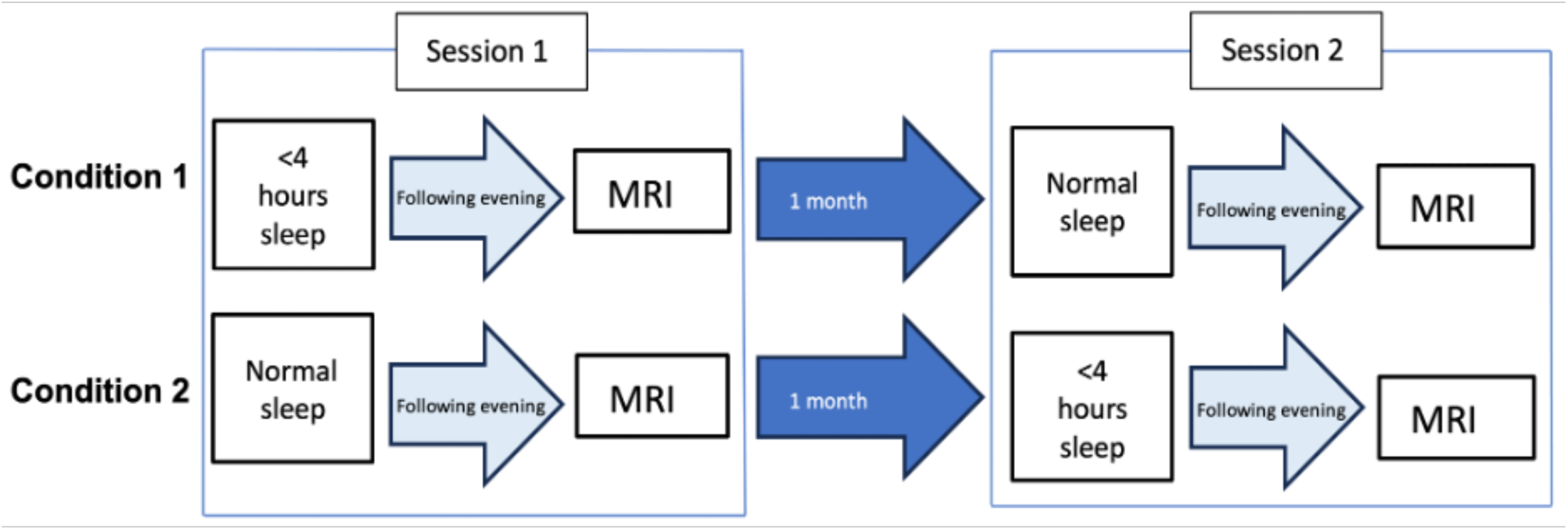
Experimental Protocol. Participants were randomly assigned to one of two conditions. Condition 1 involved partial sleep deprivation at session 1, followed by a night of normal sleep a month later. Condition 2 involved a night of normal sleep followed by a night of partial sleep deprivation a month later. At both sessions, participants were monitored at night via at-home polysomnography and received an MRI the following evening.

Sleep staging was performed in the original Stockholm Sleepy Brain study using guidelines published by the American Academy of Sleep Medicine (AASM), and included electroencephalography (EEG), electrooculography (EOG), and electromyography (EMG). The scoring was released as part of the public Stockholm Sleepy Brain 1 data release. Here we focus on total sleep time (TST), sleep efficiency (percent TST during time in bed), and the sleep stages. The latter refer to sleep stage N1 (EEG with theta and alpha waves only and no sleep spindles [short bursts of 12-15 Hz rhythmic waves], superficial sleep, transition from wake to sleep proper); stage N2 (EEG theta waves, sleep spindles and occasional delta [0.5-4Hz] waves, containing the major bulk of sleep); stage N3 (>20% of the EEG contains delta waves, deep sleep); and REM sleep (theta activity, no sleep spindles, rapid eye movements, low EMG tonus). All MRIs were acquired the evening following the recorded sleep. Participants were tracked with actigraphy data to ensure that they did not fall asleep between awakening and returning to the research center, and all participants showing signs of insomnia, irregular sleep patterns (out-of-circadian), and sleep apnea via their responses to the Insomnia Severity Index and Karolinska Sleep Questionnaire were excluded. Additionally, participants showing signs of depression were excluded via their Hospital Anxiety and Depression Scale scores. While the study collected data from 49 young (ages 20-30) and 41 old (ages 65-75) participants, our sample size was smaller following application of exclusion criteria. We excluded any participants who slept more than 4 hours during the partial sleep deprivation night, as well as those who had less than two hours difference in total sleep time between the night of normal sleep and the night of partial sleep deprivation. Additionally, we removed any subjects with missing imaging data, or with excessive motion in the image leading to poor PVS segmentation quality. This left us with 116 unique data points representing 60 unique subjects. Group differences in polysomnography results have been explored in-depth in a prior publication.^26^ **Table 1** presents the complete summary statistics for our cohort.

**Table 1:** Descriptive Statistics of Complete Cohort.

**A) Demographics**
|  | **Normal Sleep**<br>**(N=60)** | **Sleep Deprived**<br>**(N=56)** |
| --- | --- | --- |
| **Age Group** |  |  |
| Older (%) | 27 (45%) | 27 (48%) |
| Younger (%) | 33 (55%) | 29 (52%) |
| **Sex** |  |  |
| Female (%) | 29 (48%) | 28 (50%) |
| Male (%) | 31 (52%) | 28 (50%) |
| **BMI** | 24±3.8 | 24±3.8 |
| **Session** |  |  |
| 1 (%) | 30 (50%) | 28 (50%) |
| 2 (%) | 30 (50%) | 28 (50%) |

Notes: Participant demographics stratified by sleep deprivation condition for each relevant variable are included below. Mean and standard deviation are reported for continuous variables, while *N* as a percent of population is reported for categorical variables.

**B) Polysomnography and Brain Volumetric Measures for Normal Sleep versus Sleep Deprivation**
|  | **Normal Sleep**<br>**(N=60)** | **Sleep Deprived**<br>**(N=56)** | **F-statistic** |
| --- | --- | --- | --- |
| **TST - mins** | 391±72 | 160±24 | **519.66\*\*** |
| **SE - %** | 83±10 | 86±10 | 1.706 |
| **N1 - %** | 24±17 | 20±14 | 2.298 |
| **N1 - mins** | 90±59 | 30±19 | **53.424\*\*** |
| **N2 - %** | 40±13 | 40±13 | 0.006 |
| **N2 - mins** | 160±66 | 65±26 | **103.639\*\*** |
| **N3 - %** | 12±7.8 | 24±16 | **25.984\*\*** |
| **N3 - mins** | 51±33 | 39±26 | **4.529\*** |
| **REM - %** | 23±6.9 | 16±8.7 | **22.4\*\*** |
| **REM - mins** | 90±30 | 26±14 | **210.096\*\*** |
| **CSO PVS Volume Fraction** | 0.0075±0.0066 | 0.0078±0.006 | 0.072 |
| **BG PVS Volume Fraction** | 0.0045±0.002 | 0.0044±0.0021 | 0.003 |

Notes: TST = Total sleep time (minutes), SE = sleep efficiency (% TST during time in bed), CSO = centrum semiovale, BG = basal ganglia, PVS = perivascular space; Mean, standard deviation, and associated F-statistic for polysomnography-related variables and brain volumetric data during the night of normal sleep and sleep deprivation are reported. F-statistic reports differences between normal sleep and sleep
deprivation for each of the listed measures via ANOVA. Statistical significance markers for all tables: \* p<0.05; \*\* p<0.01

### MRI Processing and PVS Segmentation

Subjects were scanned on a GE 3T Discovery 750 MRI scanner. T1 weighted images were acquired with a slice thickness of 1mm interleaved bottom to top. Available scans were processed using the recon-all feature of FreeSurfer. White matter hyperintensity (WMH) masks were generated using T1 weighted and FLAIR images via the Sequence Adaptive Multimodal Segmentation tool^29^ and the Lesion Growth Algorithm^30^ as this was shown to improve sensitivity and specificity of PVS segmentation.^19,28^ Following this, an inverse WMH mask was created. This mask contained all white matter that was not identified as WMH via the pipeline. Only PVS falling within this mask were included in the PVS mask output. PVS were segmented using an existing technique. This technique is described in detail in prior publications,^27,28^ but was modified to only require input from a T1w image using the method from Sepehrband et al.^19,27^ The segmentation on T1w only was shown to have >90% accuracy and precision in detecting PVS.^19^ In brief, the algorithm filters the images to remove high frequency noise. Then, it utilizes a frangi-filter to identify “vesselness” probability for each voxel to guide the segmentation of the tubular PVS structures. The vesselness probability map was then applied with thresholds of 0.00002 in the white matter and 0.00015 in the basal ganglia, optimized for this dataset based on multiple expert opinions. The thresholded map was then binarized to obtain PVS voxels (**Figure 2**). Images were manually inspected to ensure that segmentations were accurate, and numerical ratings were given for approximately half of all available scans to statistically assess for any bias in image and segmentation quality between the two scan sessions (**Table S1**).

**Figure 2:**
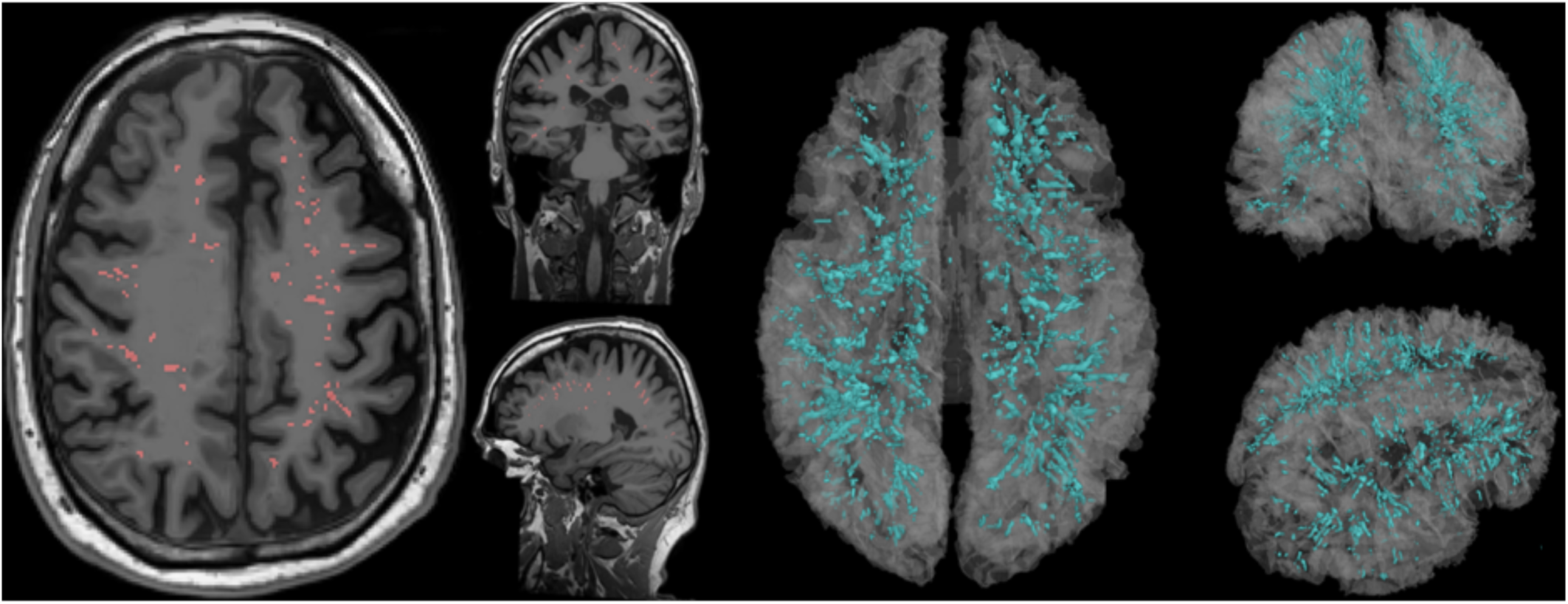
Example of 3D perivascular space segmentation technique. Visualized are the axial, coronal, and sagittal views of the segmented perivascular spaces (PVS) from one example participant in the study. At left, PVS are represented on each 2D plane (red) of the denoised T1w image. At right, 3D segmentation of PVS (blue) is overlaid on a 3D representation of the brain (transparent white).

Regional volumes in both the right and left hemisphere for each of 34 different regions within the Desikan-Killiany atlas were obtained, and volume of the PVS mask falling within each of these regions was then computed. PVS volume fractions were obtained by dividing the volume of the PVS in the region by the white matter volume of that same region. CSO PVS volume and volume fractions were obtained by adding up PVS volumes of each of the regions known to comprise the CSO (**Table S2**), and then dividing the derived total PVS volumes by the corresponding white matter volumes of these added regions, respectively. BG-PVS volume fractions were calculated by using the volume of PVS found within the basal ganglia and then dividing by the derived total basal ganglia volume using the technique described in a prior publication.^27^ As changes in the volumes of FreeSurfer generated masks have the potential to bias PVS results if brain volume segmentations change size over time, the volumes of the CSO and BG masks were checked for significant differences between scans. No significant differences were found between CSO mask volume or BG mask volume between sessions 1 and 2, nor between the scan following the night of normal sleep and the scan following sleep deprivation.

### Statistical Analysis

Paired Wilcoxon tests were conducted in R version 4.2.3.^31^ Complete information on packages used can be found in the Supplementary Material, R Packages section. Paired Wilcoxon tests were carried out to assess differences in PVS volume fraction between the nights of normal sleep and sleep deprivation. The tests were performed for each of 5 brain regions (Frontal, Occipital, Temporal, Parietal, Cingulate)^32^ in both the right and left hemispheres, resulting in a total of 10 regions.

To model the effects of sleep deprivation and time spent in each sleep stage, we utilized a linear mixed model (LMM) to account for repeated measures within subjects. Our models assessed the change in PVS volume in relation to changes in polysomnography-derived variables, including total sleep time, sleep efficiency, and both proportion and absolute duration of time spent in each stage of sleep (N1, N2, N3, REM, N2+N3). The outcome variable in all models (PVS volume fraction) was log transformed to meet model assumptions. Reported estimates and confidence intervals have been back-transformed (i.e., exponentiated) for ease of interpretability. Statistical analysis for all LMMs were performed using the *lme4* and *lmerTest* packages in R.^33,34^ Two-level LMM models were performed with repeated measures nested within subjects as a subject specific random intercept. Our outcome variables of interest were either CSO PVS volume fraction or BG-PVS volume fraction. The main predictor of interest for each model was the change in the polysomnography derived measure, including proportion of time in each sleep stage, absolute time in minutes in each sleep stage, sleep efficiency, and total sleep time. Additional variables included: sleep deprivation status, age group, sex, BMI, and MRI session number (as ordering of sleep deprivation status was counterbalanced). We included a 2-way interaction between sleep deprivation status and time spent in each sleep stage to explore whether sleep deprivation modifies the relationship between each polysomnography measure and PVS volume fraction. This approach allowed us to assess if the association between time in each stage and PVS volume fraction differed depending on participants’ sleep-deprived state. We also included four additional models without interaction effects. Two of these models assessed the main effect of sleep deprivation status (yes or no) on PVS volume fraction in the CSO and BG regions, respectively. The other two models evaluated the main effect of total sleep time on PVS volume fraction in the CSO and BG regions. Interaction effects were not included in these models, as both total sleep time and sleep deprivation status independently indicate sleep deprivation. The LMM model formulation in R format can be found in the Supplementary Material (**Formula S1**). All sleep-duration related variables (including TST, N1, N2, N3, REM, N2+N3, and sleep efficiency) were standardized (i.e., mean centered and divided by their standard deviation) to alleviate issues with model multicollinearity.

Detailed model assumption checks were conducted to ensure the use of optimal models. Sensitivity analyses were performed via paired sample t-tests to examine differences in PVS segmentation and image quality both at session 1 and session 2 and for the sleep deprived versus non-sleep deprived states to ensure that image quality and PVS segmentation quality did not significantly influence our results. Multiple comparison corrections were performed using FDR correction.

Post-hoc analyses examining a combined N2+N3 metric (for both proportion of total sleep time and time in minutes) were carried out to determine if change in N2+N3 could also be related to CSO and basal ganglia PVS volume fraction, since delta waves can be found in both sleep stages. All other covariates remained the same.

## Results

### Difference in PVS volume fraction between night of normal sleep and night of sleep deprivation

Paired Wilcoxon signed-rank tests were carried out to determine differences in PVS volume fraction between the sleep deprived and non-sleep deprived conditions. When all subjects were looked at as a complete cohort, we did not find any significant difference in PVS volume fraction between sleep deprivation and normal sleep in each of the 5 regions across each hemisphere. Complete results can be visualized in **Table S3**.

### Effect of sleep deprivation status on centrum semiovale & basal ganglia perivascular space volume fraction

#### Sleep Deprivation Status & Total Sleep Time

We utilized an LMM model to assess the overall effects of sleep deprivation on CSO and BG PVS volume fraction in the absence of interaction terms. There were no significant main effects of sleep deprivation in the CSO (*β*=1.05, p=0.781) or in the BG. (*β*=0.974, p=0.781). We performed additional models using total sleep time (TST) in place of the sleep deprivation status variable and found no main effects of total sleep time (TST) on either the CSO (*β*=0.970, p=0.781) or BG PVS volume fraction (*β*=1.02, p=0.781) (**Table S4**).

#### Sleep Efficiency

We did not observe any significant interaction effects between sleep deprivation status and sleep efficiency on PVS volume fraction (**Table S4**). The effect of sleep efficiency, defined as total time spent sleeping divided by total time in bed, was not significantly associated PVS volume fraction in either the CSO (*β*=1.02, p=0.783) or basal ganglia (*β*=1.02, p=0.783) when sleep deprivation status was at its reference level (no sleep deprivation). Similarly, the effect of sleep deprivation on PVS volume fraction, when sleep efficiency was at its average value (M=84.5%), was not significant in the CSO (*β*=1.04, p=0.781) or basal ganglia (*β*=0.969, p=0.781).

### Influence of change in both proportion and absolute time in each sleep stage on centrum semiovale PVS volume fraction

For each model, we included an interaction between sleep deprivation status and the sleep variable of interest (time in stage as either percentage of total sleep or absolute minutes) (**Figure 3**).

**Figure 3:**
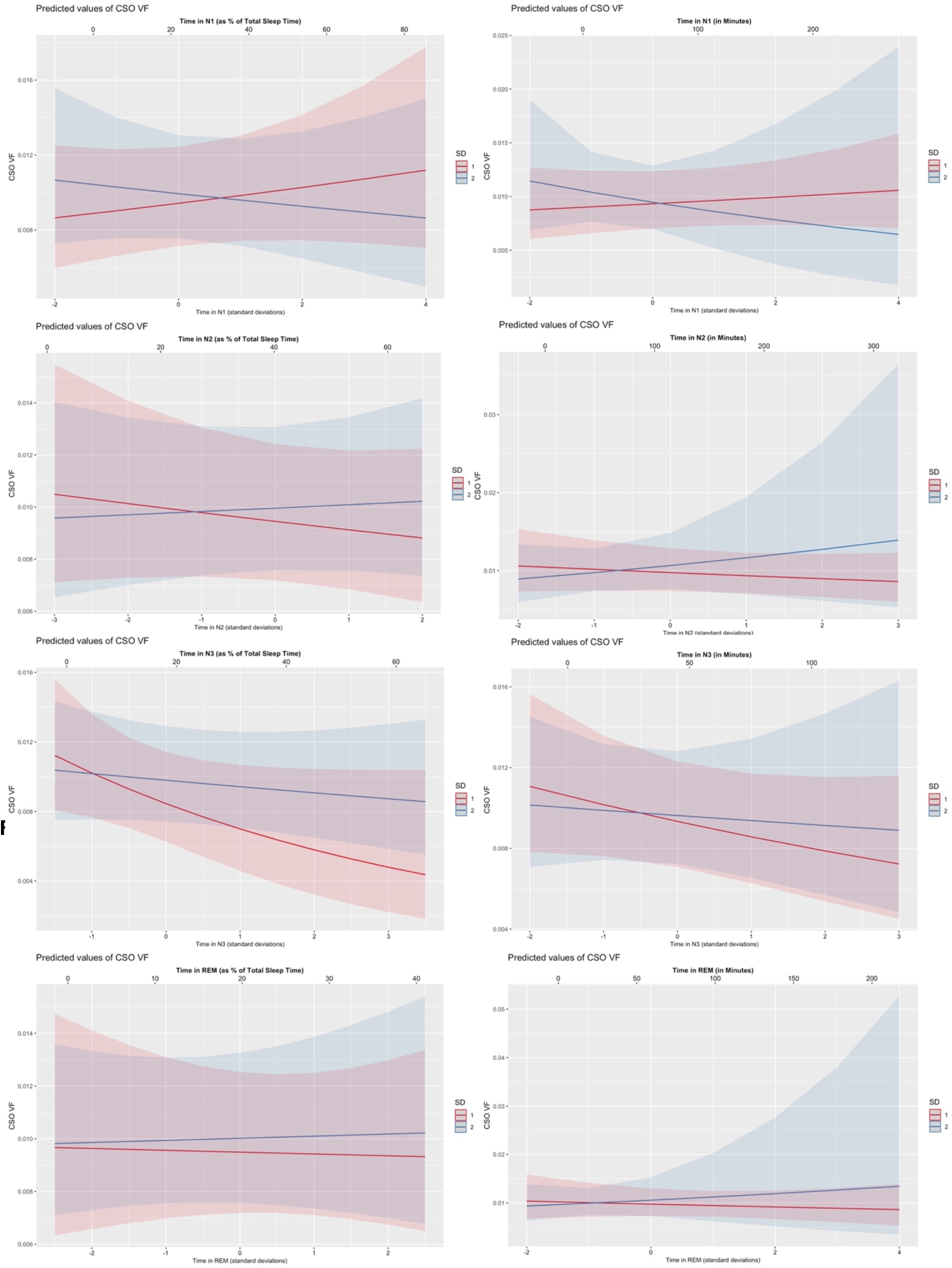
Relationship between time in sleep stage and centrum semiovale perivascular space volume fraction modified by sleep deprivation status. First column represents the relationship between percentage of time spent in each stage of sleep on centrum semiovale (CSO) perivascular space (PVS) volume fraction (VF). Trajectories were different between non-sleep deprived and sleep deprived. Second column represents the relationship between the absolute duration in minutes spent in each stage and corresponding CSO PVS VF. X-axis below each plot shows standardized values (i.e., standard deviations from the mean) of time in each respective sleep stage. X-axis above each plot shows raw values for the time spent in each sleep stage. SD = sleep deprivation status (1 = not sleep deprived [red], 2 = sleep deprived [blue]).

**Figure 4:**
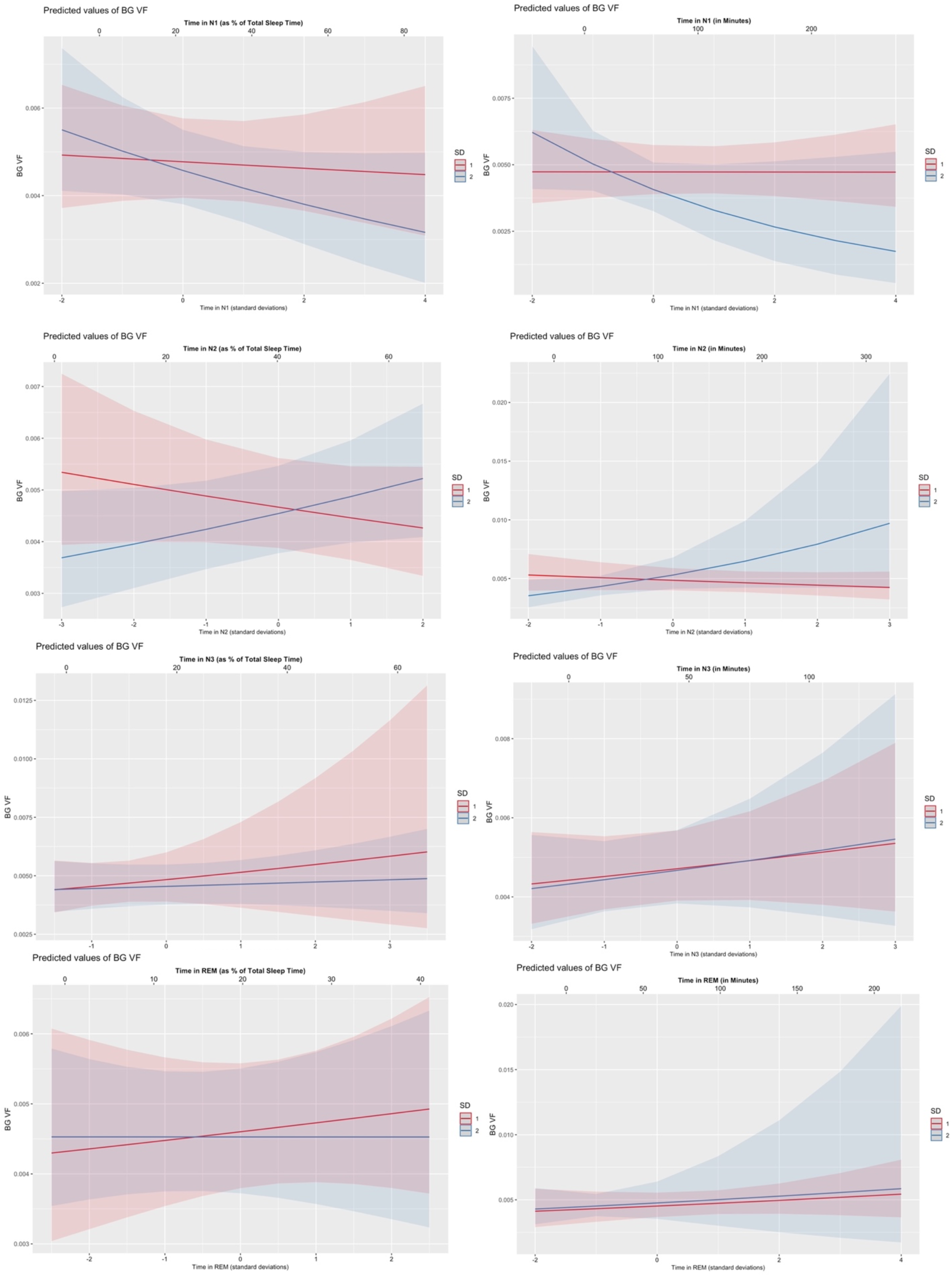
Relationship between time in sleep stage and basal ganglia perivascular space volume fraction modified by sleep deprivation status. First column represents the relationship between percentage of time spent in each stage of sleep on basal ganglia (BG) perivascular space (PVS) volume fraction (VF). Trajectories were different between non-sleep deprived and sleep deprived. Second column represents the relationship between the absolute duration in minutes spent in each stage and corresponding BG PVS VF. X-axis below each plot shows standardized values (i.e., standard deviations from the mean) of time in each respective sleep stage X-axis above each plot shows raw values for the time spent in each sleep stage. SD = sleep deprivation status (1 = not sleep deprived [red], 2 = sleep deprived [blue]).

#### N1

There was no significant interaction between sleep deprivation status and percent time in N1 on PVS volume fraction (**Table 2**). The effect of change in proportion of time spent in N1 sleep was not significantly associated with PVS volume fraction in the CSO when sleep deprivation was at its reference level (no sleep deprivation) (*β*=1.04, p=0.665). Additionally, there was no significant effect of sleep deprivation when N1% was at its reference level (M=21.93%) (*β*= 1.05, p=0.665). For the model examining absolute time in minutes in N1, there was no significant interaction between N1 in minutes and sleep deprivation status (*β*=0.881, p=0.665). Additionally, there were no significant effect of change in absolute time in minutes in N1 when sleep deprivation was at the reference level of no sleep deprivation (*β*=1.03, p=0.665), nor any significant effects of sleep deprivation (*β*=1.01, p=0.897) when absolute time in minutes of N1 was at its reference level (M=60.71 minutes).

**Table 2:** Results from linear mixed models showing effects of change in sleep stage (either in minutes or as a percentage of total sleep) on centrum semiovale PVS volume fraction.

| SLEEP MEASURE | ESTIMATE | CI | P-VALUE [uncorrected] |
| --- | --- | --- | --- |
| **N1, %** | 1.04 | 0.941 – 1.16 | 0.665 [0.412] |
| Sleep Deprived | 1.05 | 0.960 – 1.16 | 0.665 [0.261] |
| N1, % x Sleep Deprived | 0.925 | 0.838 – 1.02 | 0.665 [0.122] |
| **N2, %** | 0.966 | 0.881 – 1.06 | 0.665 [0.453] |
| Sleep Deprived | 1.05 | 0.962 – 1.15 | 0.665 [0.252] |
| N2, % x Sleep Deprived | 1.05 | 0.941 – 1.17 | 0.665 [0.380] |
| **N3, %** | 0.828 | 0.677 – 1.01 | 0.528 [0.066] |
| Sleep Deprived | 1.16 | 0.999 – 1.34 | **0.528 [0.051]** |
| N3, % x Sleep Deprived | 1.16 | 1.00 – 1.35 | **0.528 [0.051]** |
| **REM, %** | 0.993 | 0.888 – 1.11 | 0.897 [0.897] |
| Sleep Deprived | 1.06 | 0.943 – 1.18 | 0.665 [0.344] |
| REM, % x Sleep Deprived | 1.02 | 0.894 – 1.15 | 0.884 [0.810] |
| **N1, MINS** | 1.03 | 0.942 – 1.13 | 0.665 [0.496] |
| Sleep Deprived | 1.01 | 0.817 – 1.26 | 0.897 [0.896] |
| N1, mins x Sleep Deprived | 0.881 | 0.687 – 1.13 | 0.665 [0.314] |
| **N2, MINS** | 0.959 | 0.870 – 1.06 | 0.665 [0.393] |
| Sleep Deprived | 1.09 | 0.875 – 1.36 | 0.665 [0.430] |
| N2, mins x Sleep Deprived | 1.14 | 0.897 – 1.45 | 0.665 [0.279] |
| **N3, MINS** | 0.918 | 0.815 – 1.03 | 0.665 [0.159] |
| Sleep Deprived | 1.03 | 0.932 – 1.14 | 0.686 [0.544] |
| N3, mins x Sleep Deprived | 1.06 | 0.958 – 1.17 | 0.665 [0.253] |
| **REM, MINS** | 0.970 | 0.860 – 1.09 | 0.698 [0.611] |
| Sleep Deprived | 1.08 | 0.818 – 1.43 | 0.686 [0.572] |
| REM, mins x Sleep Deprived | 1.10 | 0.838 – 1.43 | 0.665 [0.499] |

Notes: Parameter estimates, confidence intervals, and both corrected and uncorrected p-values for each model are listed. Statistical significance markers: \* p<0.05; \*\* p<0.01; \*\*\* p<0.001

#### N2

We did not observe any significant interaction effect between change in time spent in N2 and sleep deprivation status (**Table 2**). No significant effect of change in either percent time (*β*=0.966, p=0.665) or total time in minutes in N2 (*β*=0.959, p=0.665) was found to be associated with CSO PVS volume fraction when sleep deprivation status was at the reference level (no sleep deprivation). We also did not observe any significant effect of sleep deprivation in these models, when N2% (M=40.19%) or N2 mins (M=114.28 minutes) were at their respective reference levels.

#### N3

We noted a borderline result showing a possible interaction effect between sleep deprivation status and change in percent time spent in N3 on CSO PVS volume fraction (*β*=1.16, p=0.528 [uncorrected p=0.051]). The cross-over point (referring to the proportion of time in N3 at which both groups showed equivalent effects on CSO PVS volume fraction) occurred at -0.98 standard deviations below the mean (equivalent to 4.81% N3, given M=18.09%, SD=13.5). Overall, the non-sleep deprived group showing faster decreases in CSO PVS volume fraction with increasing percentage of time spent in N3 compared to the sleep deprived group. However, this result did not survive multiple comparisons corrections. No significant interaction effect was seen for absolute time spent in N3 (in minutes) and sleep deprivation status on CSO PVS volume fraction (**Table 2**).

#### REM

No significant interaction effects were observed between REM sleep (both in minutes and as % of TST) and sleep deprivation status on CSO PVS volume fraction (**Table 2**). Change in percent REM sleep was not significantly associated with CSO PVS volume fraction when sleep deprivation was at the reference level of no sleep deprivation (*β*=0.993, p=0.897). Change in time spent in REM minutes was also not significantly associated with CSO PVS volume fraction (*β*=0.970, p=0.698) at the reference level (no sleep deprivation).

### Influence of change in both proportion and absolute time in each sleep stage on basal ganglia PVS

#### N1

We did not observe any significant interaction effect between time in N1 sleep and sleep deprivation status in either of the N1 models for BG (**Table 3**). Additionally, we did not see any significant effect of change in N1% (*β*=0.984, p=0.911) or duration in minutes of N1 sleep (*β*=1.00, p=0.992) on basal ganglia PVS volume fraction at the reference level (no sleep deprivation).

**Table 3:** Results from linear mixed models showing effects of change in sleep stage (either in minutes or as a percentage of total sleep) on basal ganglia PVS volume fraction.

| SLEEP MEASURE | ESTIMATE | CI | P-VALUE [uncorrected] |
| --- | --- | --- | --- |
| **N1, %** | 0.984 | 0.900 – 1.08 | 0.911 [0.725] |
| Sleep Deprived | 0.958 | 0.876 – 1.05 | 0.856 [0.348] |
| N1, % x Sleep Deprived | 0.926 | 0.842 – 1.02 | 0.586 [0.114] |
| **N2, %** | 0.956 | 0.882 – 1.04 | 0.856 [0.270] |
| Sleep Deprived | 0.974 | 0.895 – 1.06 | 0.884 [0.529] |
| N2, % x Sleep Deprived | 1.12 | 1.02 – 1.24 | **0.300 [0.024]** |
| **N3, %** | 1.06 | 0.884 – 1.28 | 0.884 [0.504] |
| Sleep Deprived | 0.940 | 0.817 – 1.08 | 0.856 [0.381] |
| N3, % x Sleep Deprived | 0.959 | 0.827 – 1.11 | 0.884 [0.568] |
| **REM, %** | 1.03 | 0.930 – 1.14 | 0.884 [0.589] |
| Sleep Deprived | 0.984 | 0.885 – 1.09 | 0.911 [0.759] |
| REM, % x Sleep Deprived | 0.973 | 0.864 – 1.10 | 0.909 [0.647] |
| **N1, MINS** | 1.00 | 0.922 – 1.08 | 0.992 [0.992] |
| Sleep Deprived | 0.860 | 0.710 – 1.04 | 0.586 [0.122] |
| N1, mins x Sleep Deprived | 0.809 | 0.645 – 1.02 | 0.536 [0.067] |
| **N2, MINS** | 0.956 | 0.878 – 1.04 | 0.856 [0.301] |
| Sleep Deprived | 1.09 | 0.898 – 1.33 | 0.856 [0.376] |
| N2, mins x Sleep Deprived | 1.28 | 1.03 – 1.59 | **0.300 [0.025]** |
| **N3, MINS** | 1.04 | 0.938 – 1.16 | 0.856 [0.428] |
| Sleep Deprived | 0.991 | 0.898 – 1.09 | 0.938 [0.860] |
| N3, mins x Sleep Deprived | 1.01 | 0.913 – 1.12 | 0.938 [0.852] |
| **REM, MINS** | 1.05 | 0.940 – 1.17 | 0.856 [0.397] |
| Sleep Deprived | 1.05 | 0.821 – 1.35 | 0.909 [0.682] |
| REM, mins x Sleep Deprived | 1.01 | 0.785 – 1.29 | 0.992 [0.963] |

Parameter estimates, confidence intervals, and both corrected and uncorrected p-values for each model are listed. Statistical significance markers: \* p<0.05; \*\* p<0.01; \*\*\* p<0.001

#### N2

We observed interaction effects in each of our N2 models (**Table 3**). In the percent N2 model, an increasing proportion of time spent in N2 was associated with decreases in BG PVS volume fraction in the non-sleep deprived group, while the sleep deprived group showed the opposite effect, with increases in BG PVS volume fraction as the proportion of time in N2 increased (*β*=1.12, p=0.300 [uncorrected p=0.024]). Cross-over, or the point at which effects in each group were equivalent in the percent N2 model, occurred at 0.23 standard deviations above the mean (equivalent to 43.17% N2, given M=40.19%, SD=12.97). In the N2 minutes model, we observed a similar interaction effect, with the non-sleep deprived group showing decreases in BG PVS volume fraction as N2 increased, while the sleep deprived group showed increases in N2 associated with increases in BG PVS volume (*β*=1.28, p=0.300 [uncorrected p=0.025]). In this model, cross-over occurred at -0.35 standard deviations below the mean (equivalent to 89.98 minutes of N2, given M=114.28 minutes, SD=69.44 minutes).

#### N3

We did not observe any significant interaction effects between time in N3 (both minutes and percent of TST) and sleep deprivation (**Table 3**). No significant effects of percent N3 (*β*=1.06, p=0.884) or minutes N3 (*β*=1.04, p=0.856) were observed when sleep deprivation was at the reference level (no sleep deprivation). Additionally, there were no effects of sleep deprivation alone when time in N3 was at the reference level for percent N3 (M=18.09%) and N3 minutes (M=45.04 minutes).

#### REM

No interactions between REM minutes or percent REM and sleep deprivation were observed (**Table 3**). We also did not find any significant effect of time in REM sleep in minutes (*β*=1.05, p=0.856) or as a percentage of total sleep (*β*=1.03, p=0.884) when sleep deprivation was at the reference level (no sleep deprivation). Additionally, we did not find any main effect of sleep deprivation when time in REM was at the reference levels for percent REM (M=19.74%) and REM minutes (M=59.03 minutes).

### Post-hoc analyses

#### N2+N3

Duration and proportion of time spent in stages N2 and N3 were combined in a post-hoc analysis, as both stages include some delta activity. There was no significant interaction effect of time in the combined N2+N3 metric (both in minutes and as percentage of total sleep time) and sleep deprivation status (**Table S5**). Combined change in proportion of time spent in N2+N3 was not significantly associated with CSO PVS volume fraction at the reference level of non-sleep deprived (*β*=0.936, p=0.547). Time spent in N2+N3 in minutes was also not significantly associated with CSO (*β*=0.965, p=0.638) or basal ganglia PVS volume fraction (*β*=1.00, p=0.923) at the reference level (non-sleep deprived). There was no main effect of sleep deprivation in both CSO and BG models at either of their respective reference levels for percent N2+N3 (M=58.28%) or time spent in N2+N3 in minutes (N=159.31).

## Discussion

Our study aimed to elucidate within-subject differences in PVS volume fraction resulting from participation in a late-night short sleep deprivation study. Overall, we found no significant differences in PVS volume fraction between a night of normal sleep and a night of partial sleep deprivation. The lack of findings indicates that partial sleep deprivation may not be sufficient for assessing effects of sleep loss on PVS changes. It could also be possible that acute changes in sleep are not conducive to observing such changes, and that sustained poor sleep may be required for any effects of poor sleep on PVS to be observed. Despite the lack of significant findings, this illustrates new areas of focus for future investigators. Additionally, while none of our results survived multiple comparisons correction, we did note a few interesting trends that were significant before corrections which will be expanded upon in this section. It is possible that our large number of tests, coupled with only mild effects of sleep deprivation due to the nature of the late-night short-sleep experiment, led to missed associations, but future work will be needed to determine with certainty if this is the case.

We suspected that deficits in sleep duration, particularly deep sleep, would lead to increases in PVS volume fraction. Our results partially support this notion as we found an initial marginal interaction between proportion of time spent in N3 and sleep deprivation status on the CSO PVS volume fraction. This trend seems to show that those who are not sleep deprived may experience decreases in CSO PVS volume fraction as they receive more N3 sleep, while the effect is less pronounced in the sleep deprived state. Despite sleep-deprived individuals spending relatively more of the night in N3 (24±16% of TST vs. 12±7.8% of TST for non-sleep deprived individuals), this shows that the increased time in N3 during late-night short-sleep may not be enough to compensate for a full night of sleep. These findings are somewhat consistent with previous work, such as the recent findings by Baril et al,^14^ which found associations between higher absolute time in N1 and lower absolute time in N3 with heightened PVS burden. Another study by Yang et al,^22^ found a significant association between poor sleep quality, measured via the Pittsburgh Sleep Quality Index (PSQI), and enlarged PVS in both the centrum semiovale and basal ganglia. Opel et al,^35^ found an association between lower overall total sleep time and greater enlarged PVS burden. Since it’s possible that the effect we observed was only mildly detectable due to our small sample size, future studies should investigate larger cohorts and include more lengthy sleep deprivation experiments that span several days and include several follow-up MRI acquisitions.

Additionally, we observed an initial interaction effect of both N2 (percent and minutes) and sleep deprivation status in the basal ganglia. For both cases, it appears that increases in N2 during sleep deprivation are linked to higher basal ganglia PVS volume fraction, while increases in N2 during normal sleep are associated with decreased PVS volume fraction. This again upholds the idea that regular sleep mechanics are impaired in late-night short-sleep and increases in time in the deeper stages (as N2 contains some delta activity) could result in larger PVS volume fraction in those who are sleep deprived.

One potential hypothesis could be that these contradictory effects exist as a compensatory mechanism for lost sleep. It is known that sleep deprivation will affect sleep architecture, so it is possible that sleep deprivation could also alter fluid flow patterns to enhance or better optimize the clearance given the limitation of impaired sleep. Future studies are needed to look at longer term effects of late-night short sleep deprivation, with multiple follow-ups and extensive sleep tracking over time necessary to establish any clear links in humans.

### Late Night Short Sleep and Generalizability to the Population at Large

This study’s focus on late-night short sleep allows the results to be more applicable to a larger population. Most individuals with sleep impairment in the clinical setting do not undergo absolute lack of sleep, but rather difficulty falling and staying asleep. This problem is especially apparent in older individuals, where nearly 13 percent of individuals above age 65 years report either moderate or severe sleep insufficiency.^36^ Late-night short sleep experiments represent a method of investigating shorter sleep times on the brain, rather than absolute sleep deprivation which is unlikely to occur regularly.

Prior literature investigating the role of the brain’s PVS system during sleep indicate that waste clearance occurs most readily during deep sleep, measured via influx of concentrations of molecular tracers from the CSF of the cisterna magna, along the periarterial space, and through the parenchyma.^37^ The prior literature using tracers does not consider size of PVS on structural MRI imaging. PVS observed on MRI are currently hypothesized to represent influx of CSF into the parenchyma,^38^ becoming enlarged as a result of impaired clearance. The growth of the PVS likely represent stagnant or slowed fluid flow, though future studies should aim to combine both contrast-enhanced or functional imaging with structural imaging to see if there is a link between structural changes in the PVS over a short period of time, such as one night of partial sleep deprivation, and clearance processes. Studies should also aim to examine effects in old and young age groups separately, as it is possible mechanisms between age groups may be slightly different due to physiological changes that occur during aging.

Something to consider about our study is that it involved a relatively homogenous group of individuals who may not be as affected by sleep as a risk factor for brain health compared to other populations. While these participants may represent a largely educated population with few health conditions and generally good sleep, examining the effects of short-term sleep deprivation in historically underrepresented groups or those who face systemic barriers to quality sleep may yield different results.^39^ This points to the importance of including such groups in future studies.

### Strengths and Limitations

Our work has several notable strengths and weaknesses. Use of polysomnography is considered the gold standard in sleep research and allows for quantification of changes in sleep measures with variables of interest. Participants in this study had polysomnography done in-home, so they did not require any acclimation to sleep in an unfamiliar environment, leading to more realistic recorded results. Having MRI done at two timepoints, one after a normal night of sleep and another after a night of partial sleep deprivation, allowed for thorough examination of the effects of partial sleep deprivation. Additionally, our automated technique that allowed for quantification of both global CSO PVS and regional PVS volumes permitted us to determine volumetric changes in PVS, as opposed to measures that are based on visual scoring alone. Quantification of PVS changes at the voxel level allows for more subtle detection of changes. Lastly, our statistical analysis allowed for the investigation of numerous potential associations between PVS change and sleep deprivation metrics, ranging from changes in proportion of time spent in each stage to simple effects of sleep deprivation alone. This allowed us to get a more in-depth look at what exact components of sleep deprivation (such as changes in proportion of N2) are contributing to changes in the PVS volume.

Another strength of our study is that we investigated changes in CSO and basal ganglia PVS volume fraction using a linear mixed-effects model, which can account for repeated observations within subjects while including additional information as covariates. In these cases, we found that it was the changes in proportion of time spent in each stage of sleep, rather than whether someone was simply sleep deprived, that related to differences in PVS volume fraction. This finding has also been previously supported in the literature.^14^ The design of the late-night short-sleep (LNSS) experiment, involving delayed bedtime and less than four hours of sleep, can also be viewed as a unique component of our study. Since LNSS typically lead to reductions in the amount of N1, N2, and REM but relatively less change in absolute N3 duration due to the emergence of compensatory homeostatic mechanisms that follow sleep loss, our experiment is able to address additional questions regarding receiving varying amounts of light sleep^40,41^ In LNSS experiments, increased sleep pressure accumulates due to the delayed bedtime, creating an earlier transition to N3 sleep. Thus, there is a homeostatic response in LNSS experiments resulting in sufficient absolute N3 minutes despite the sleep deprivation. It is likely that a total sleep deprivation protocol would be necessary to obtain more notable differences in absolute minutes in N3 between sleep deprived and non-sleep deprived states. Our study allows for examination of other stages of sleep that may contribute to PVS changes (particularly N2, as we saw with our basal ganglia results) while relatively normal amounts of N3 are maintained. However, it’s important to note that we saw sleep deprived individuals spending more time in N3 (as a percentage of their total sleep time) compared to their night of normal sleep, which may have also contributed to our decreased ability to find significant changes in PVS volume fraction following only a partial sleep deprivation protocol.

Despite our studies strengths, there are still notable limitations. The in-home polysomnography recording does not exclude potential issues with recording that could have occurred in this setting, such as participant failure to follow protocol directions. Additionally, some participants failed to follow the sleep deprivation instructions, sleeping more than the allotted time during the experimental condition. We attempted to remedy this by simply excluding participants who slept 4 hours or more from our analysis.

The imaging data used in this study were T1w images with 1mm resolution, which is less sensitive in detecting PVS compared to methods using T1w divided by T2w, also known as the enhanced PVS contrast.^27^ Ultra-high field imaging done at 7T could also improve our ability to detect subtle changes in PVS when using T1w images alone.^42^

Data collection for this study took place over the course of several months in Sweden, where summers are long, and winters are short. A recent publication investigated the effects of long summer and short winter days on circadian timing.^43^ Our study was within subject, and participants were scanned approximately one month apart, so any drastic changes in circadian timing seem unlikely, although they cannot be ruled out. Future studies should look to include seasonality as a covariate for any potential effects of circadian disruption. However, a strength of the study design involved scanning participants in the evening following the sleep deprivation condition, so influence of daily circadian changes in the brain waste clearance process did not influence our result.^44^ However, studies looking to address daily changes in PVS size due to the effects of circadian timing could aim to quantify changes in PVS based on time of day in one individual, with several imaging sessions being acquired within one day for this investigation.

Lastly, our sample size was quite small and therefore some effects may have been missed. Our small sample size resulted in very little variance in our data when fitting our LMM models, further potentially contributing to missed effects. Future studies should aim to include several hundreds or thousands of participants.

### Conclusions

Our study represents the first look at within-subject changes in PVS volume fraction in relation to a sleep deprivation experiment. The results exhibit a degree of alignment with prior literature showing that increases in N3 and decreases in N2 proportion are associated with decreased PVS volume fraction. None of our results survived multiple comparisons correction, so we are limited in conclusions we can draw about the observed effects. We suspect that we were simply limited in detecting such effects due to small sample size. Future research is needed to determine more definitive links between sleep deprivation and enlargement of PVS. This would allow sleep practitioners to better assess effects of sleep related changes on PVS alongside any potential associated risks of such enlargement.

## Acknowledgements

The research reported in this publication was supported by the National Institute of Mental Health (RF1MH123223) and National Institute on Aging of the NIH under Award Numbers R44AG087855-01, R41AG073024, and R01AG070825. The content is solely the responsibility of the authors and does not necessarily represent the official views of the NIH. The research was also supported by Riksbankens Jubilemumsfond (grant no. P15-0310:1), by the Fredrik and Ingrid Thuring Foundation (grant no. 2015-00170) and by the Rut and Ingrid Wolff Memorial Foundation (grants no. 2019-01734 and 2020-01412).

## Disclosure Statements

### Financial Disclosure

JC is receiving salary from NeuroScope Inc.

### Non-financial Disclosure

The perivascular space mapping technology is part of a patent owned by Jeiran Choupan, with no financial interest/conflict.

## Data Availability Statement

Data used in this manuscript are available in the OpenNeuro online repository [https://openneuro.org] via Accession Number ds000201 [https://openneuro.org/datasets/ds000201/versions/00004].

## Notes

https://openneuro.org/datasets/ds000201/versions/00004

### Competing Interest Statement

Financial Disclosure: Jeiran Choupan is receiving salary from NeuroScope Inc.
Non-financial Disclosure: The perivascular space mapping technology is part of a patent owned by Jeiran Choupan, with no financial interest/conflict.

