## Supplementary Material for "Effects of one-night partial sleep deprivation on perivascular space volume fraction: Findings from the Stockholm Sleepy Brain Study"

**TABLES**

**Table S1: Manual Inspection and Rating Information for Subset of Participants –** Manual ratings were performed for scans from session 1 and session 2 for a subset of participants, representing approximately 45% of all available scans. T1w images were rated on a scale of 1 (best) to 3 (worst) quality based on presence of image artifacts, motion, or image intensity issues. Automated perivascular space segmentations were inspected and rated on the same scale of 1 (best) to 3 (worst) and were evaluated poorly if there was an overabundance of false positive segmentations and segmentation of structures known to be image artifacts or other pathologies (such as white matter hyperintensities).

**Subject	Session 1 Scan	Session 1 PVS	Session 2 Scan	Session 2 PVS**

| **sub-9003** | 1.5 | 2 | **1.5** | **1** |
| --- | --- | --- | --- | --- |
| **sub-9004** | 2 | 2.5 | 1.5 | 2 |
| **sub-9009** | 2 | 2 | 2 | 2 |
| **sub-9011** | 2.5 | 2 | 1.5 | 2.5 |
| **sub-9014** | 1.5 | 2 | 1.5 | 1.5 |
| **sub-9017** | 1 | 1.5 | 1 | 1.5 |
| **sub-9019** | 1 | 1 | 1 | 1 |
| **sub-9020** | 1.5 | 1.5 | 1 | 1.5 |
| **sub-9026** | 1.5 | 2 | 1.5 | 2 |
| **sub-9028** | 2 | 1.5 | 1.5 | 1.5 |
| **sub-9032** | 2 | 2 | 1 | 1.5 |
| **sub-9033** | 2 | 2 | 2 | 2 |
| **sub-9034** | 2 | 1.5 | 1.5 | 1.5 |
| **sub-9037** | 1 | 2 | 2 | 1 |
| **sub-9038** | 1 | 1 | 2 | 1.5 |
| **sub-9039** | 1 | 2 | 2 | 2 |
| **sub-9042** | 1 | 1.5 | 1 | 1 |
| **sub-9045** | 1 | 2 | 1.5 | 1.5 |
| **sub-9046** | 1 | 1 | 1.5 | 1.5 |
| **sub-9049** | 1.5 | 1.5 | 1.5 | 2 |
| **sub-9055** | 1.5 | 1 | 1 | 1 |
| **sub-9058** | 1 | 1 | 1 | 1.5 |
| **sub-9064** | 1.5 | 1.5 | 1.5 | 2 |
| **sub-9067** | 1.5 | 1.5 | 1 | 1.5 |
| **sub-9068** | 1 | 1.5 | 1.5 | 1.5 |
| **sub-9071** | 1 | 1.5 | 1 | 1.5 |
| **sub-9072** | 1 | 1 | 1 | 1.5 |
| **sub-9073** | 2 | 1.5 | 2.5 | 2 |
| **sub-9074** | 1 | 1 | 1 | 1 |
| **sub-9082** | 2 | 2 | 1 | 1 |
| **sub-9085** | 1 | 1.5 | 1 | 1.5 |
| **sub-9086** | 1.5 | 1.5 | 1.5 | 2 |
| **sub-9090** | 1.5 | 1.5 | 1.5 | 2 |
| **sub-9091** | 1.5 | 1.5 | 1 | 1.5 |
| **sub-9092** | 1 | 1 | 1 | 1.5 |
| **sub-9094** | 1 | 1 | 2.5 | 2.5 |
| **sub-9096** | 1 | 1 | 1 | 1.5 |

**Table S2: Regions Comprising the Centrum Semiovale –** The following regions correspond to FreeSurfer identified regions of interest (ROI) in both the left and right hemispheric white matter. We used the volume of PVS in all regions of this list added together as our centrum semiovale ROI. Centrum Semiovale perivascular space (PVS) volume fraction was calculated by adding the PVS volumes from each of these regions together, then dividing by the total white matter volume of these regions combined.

**Centrum Semiovale Regions**

| Caudal Middle Frontal | Precentral |
| --- | --- |
| Inferior Parietal | Rostral Middle Frontal |
| Pars Opercularis | Superior Frontal |
| Pars Orbitalis | Superior Parietal |
| Pars Triangularis | Supramarginal |
| Postcentral |  |

**Table S3: Results of Wilcoxon Signed Rank Tests (Complete Cohort)** – Results from paired sample Wilcoxon tests examining perivascular space volume differences between the night of normal sleep and sleep deprivation for 5 different brain regions. None of these results survived FDR multiple comparison corrections.

**LEFT HEMISPHERE	RIGHT HEMISPHERE**

**P-Value	P-Value**

| FRONTAL | 0.897 | 0.897 |
| --- | --- | --- |
| OCCIPITAL | 0.897 | 0.913 |
| PARIETAL | 0.984 | 0.897 |
| TEMPORAL | 0.984 | 0.950 |
| CINGULATE | 0.897 | 0.897 |

**Table S4: Results from linear mixed models showing effects of change in sleep deprivation, total sleep time, and sleep efficiency on centrum semiovale and basal ganglia PVS volume fraction.** P values have been adjusted for multiple comparisons using FDR correction. Main effect estimates of total sleep time (TST) and sleep efficiency for each model represent the effect when the participant is not sleep-deprived (SD=1). Effect estimates for sleep efficiency refer to the estimate obtained when the Sleep Efficiency (SE) value is at the reference level (M=84.5%).

| **CENTRUM SEMIOVALE** | **ESTIMATE** | **CI** | **P-VALUE** |
| --- | --- | --- | --- |
| SLEEP DEPRIVATION | 1.05 | 0.962 – 1.16 | 0.781 |
| TOTAL SLEEP TIME (TST) | 0.970 | 0.924 – 1.02 | 0.781 |
| SLEEP EFFICIENCY (SE) | 1.02 | 0.915 – 1.15 | 0.783 |
| Sleep Deprived | 1.04 | 0.949 – 1.15 | 0.781 |
| Sleep Deprived x Sleep Efficiency | 1.03 | 0.935 – 1.14 | 0.781 |
| **BASAL GANGLIA** | **ESTIMATE** | **CI** | **P-VALUE** |
| SLEEP DEPRIVATION | 0.974 | 0.891 – 1.06 | 0.781 |
| TOTAL SLEEP TIME (TST) | 1.02 | 0.969 – 1.06 | 0.781 |
| SLEEP EFFICIENCY (SE) | 1.02 | 0.924 – 1.12 | 0.783 |
| Sleep Deprived | 0.969 | 0.884 – 1.06 | 0.781 |
| Sleep Deprived x Sleep Efficiency | 1.00 | 0.910 – 1.10 | 0.965 |

**Table S5: Post-hoc Tests – N2+N3 Analysis: Results from linear mixed models showing effects of change in N2+N3 percent and N2+N3 minutes on centrum semiovale and basal ganglia PVS volume fraction.** P values have been adjusted for multiple comparisons using FDR correction. Effect estimates of N2+N3% and N2+N3 minutes refer to the effect when the participant is not sleep-deprived (SD=1). Effect estimates of Sleep Deprived refer to the estimate obtained when the N2+N3 values are at the reference level (M=58.28% for percent models; M=159.31 minutes for minute models).

| **CENTRUM SEMIOVALE** | **ESTIMATE** | **CI** | **P VALUE** |
| --- | --- | --- | --- |
| N2+N3 (%) | 0.936 | 0.823 – 1.07 | 0.547 |
| Sleep Deprived | 1.07 | 0.955 – 1.21 | 0.547 |
| Sleep Deprived x Sleep Efficiency | 1.07 | 0.969 – 1.18 | 0.547 |
| N2+N3 (Mins) | 0.965 | 0.860 – 1.08 | 0.638 |
| Sleep Deprived | 1.11 | 0.860 – 1.42 | 0.568 |
| Sleep Deprived x Sleep Efficiency | 1.16 | 0.920 – 1.45 | 0.547 |
| **BASAL GANGLIA** | **ESTIMATE** | **CI** | **P VALUE** |
| N2+N3 (%) | 1.01 | 0.904 – 1.12 | 0.923 |
| Sleep Deprived | 0.947 | 0.851 – 1.05 | 0.547 |
| Sleep Deprived x Sleep Efficiency | 1.07 | 0.973 – 1.18 | 0.547 |
| N2+N3 (Mins) | 1.00 | 0.910 – 1.11 | 0.923 |
| Sleep Deprived | 1.10 | 0.885 – 1.37 | 0.568 |
| Sleep Deprived x Sleep Efficiency | 1.20 | 0.974 – 1.48 | 0.547 |

**FORMULAS**

**Formula S1:** An example of the linear mixed effect model (LMM) using lmer command in R; changes in PVS volume in relation to the polysomnography derived sleep variable is shown below. PVS volume fraction represents the outcome variable of PVS volume fraction within either the centrum semiovale (CSO) or basal ganglia (BG). Time in sleep stage represents the total time (either in minutes or as a percentage of total sleep time) spent in each stage of sleep. These values have been standardized, so are represented in terms of standard deviations from the mean of each sleep duration variable. Sleep deprivation status is a binary variable referring to whether the participants were sleep deprived or not. Session represents the MRI session number. Age group represents the age category participants fall into (old or young). Sex represents the sex grouping variable. BMI is included as a continuous variable. The random effect of repeated measures within subject is represented by (1|Subject). The model was specified to run with REML=FALSE to specify usage of the Maximum Likelihood estimation.

lmer(log(PVS Volume Fraction) ~ Time in Sleep Stage (standardized) * Sleep Deprivation Status + Session + Age Group + Sex + BMI + (1|Subject), REML=FALSE, data=data-frame name)

**R PACKAGES**

All statistical analyses were performed in R (v4.2.3; R Core Team 2023). Additional packages and libraries used in addition to the base R packages include: *HLMdiag*^1^, *car*^2^, *emmeans*^3^, *fitdistrplus*^4^, *performance*^5^, *ggResidpanel*^6^, *sjPlot*^7^, *vtable*^8^, *remef*^9^, *sjmisc*^10^, *sjlabelled*^11^, *tidyverse*^12^, *dplyr*^13^, *plyr*^14^, *ggplot2*^15^, *scales*^16^, *corrplot*^17^, *psych*^18^, and *nlme*^19^. All main analyses were conducted using the *lme4*^20^ and *lmerTest*^21^ packages.
